# UltraPrep is a scalable, cost-effective, bead-based method for purifying cell free DNA

**DOI:** 10.1101/2020.04.03.023499

**Authors:** Christopher K. Raymond, Fenella C. Raymond, Kay Hill

## Abstract

UltraPrep is an open-source, two-step method for purification of cell-free DNA that entails extraction of total DNA followed by size-selective enrichment of the smaller fragments that are characteristic of DNA originating from fragmentation between nucleosome. The advantages of the method are that it can easily accommodate a wide range of sample input volumes, it relies on simple, magnetic bead-based technology, the yields of cfDNA are directly comparable to the most popular methods for cfDNA purification, and it dramatically reduces the cost of cfDNA isolation relative to currently available commercial methods. We provide a framework for physical and molecular quality analysis of purified cfDNA and demonstrate that the cfDNA generated by UltraPrep meets or exceeds the quality metrics of the most commonly used procedure. Our method removes high molecular weight genomic DNA that can interfere with downstream assay results, thereby addressing one of the primary concerns for preanalytical collection of blood samples.

## Introduction

Circulating cell-free DNA (cfDNA) has shown tremendous utility as an analyte in prenatal genetic analysis and in precision medicine approaches to diagnosed cancers. It holds promise to contribute to early detection of solid tumors [1,2]. Early detection tests that use cfDNA must be both highly sensitive and specific. Straightforward probability and statistics considerations indicate that this requires high input levels of cfDNA and subsequent genomic analysis that covers several thousand independent cfDNA “genome equivalents” [3,4]. In addition to individual patient testing, there is a largely unmet need for large, well-characterized, single-donor lots of normal human cfDNA that can be used for diagnostic test research, assay development, and routine proficiency qualification in clinical laboratory environments.

Plasmapheresis is a method that can be used to safely collect hundreds of milliliters of plasma from human subjects. Plasma is most often collected into vessels containing sodium citrate, a molecule that chelates divalent cations required by DNAse enzymes and thereby stabilizes extracellular DNA. Plasmapheresis samples contain significant quantities of cfDNA, and these can be used in the contexts described in the previous paragraph. At present, it is difficult to realize the potential of this cfDNA source owing to a lack of cost-effective methods for high volume cfDNA extraction and purification.

We were inspired by a recent publication promoting magnetic bead-based laboratory methods [5] to pursue an open-source approach to high-volume, reduced-cost purification of cfDNA. This proved to be a significant challenge. The most formidable obstacle was to achieve near-quantitative recovery of cfDNA fragments. Specifically, DNA binding to silica surfaces has been used as a purification method for decades [6,7], but a significant fraction of DNA is bound irreversibly [8,9]. Conditions for robust and reversible binding of cfDNA are reported here. In addition, preanalytical collection conditions often result in plasma that is a mixture of high molecular weight genomic DNA and cfDNA. By cfDNA, we mean a set of DNA fragments derived from cleavage between adjacent nucleosomes [10,11]. Since a single nucleosomal subunit is about 165 bp and cleavage between subunits can be incomplete, this results in a “ladder” of DNA fragments that are nucleosomal monomers, dimers, trimers, etc. [12]. This collection of “nucleosomal fragments” is thought to be generated by apoptosis that occurs among the cells in both normal and cancerous tissues. In contrast, high molecular weight genomic DNA that is observed in some plasma samples is thought to be largely contributed by nucleated blood cells that burst. This type of DNA can confound certain types of downstream genomic analysis such as quantitative digital PCR and amplicon-based DNA sequencing. There has been considerable effort invested in preanalytical collection methods that prevent cell lysis. Here we provide an alternative approach in which high molecular weight genomic DNA can be removed from nucleosomal fragments by bead-based partitioning.

## Materials and methods

### Ethics Statement

The ethics committees of Resolution Bioscience and of Plasma Lab International reviewed and approved of the research presented here. Written consent was obtained from healthy donors prior to sample collection, processing and characterization.

### Materials

Plasma samples were collected at PlasmaLab International (Everett, WA). Proteinase K was purchased from Gold Biotechnology (St. Louis, MO), guanidinium isothiocyanate (GITC) from Chem Impex (Wood Dale, IL), silica coated superparamagnetic beads from Spherotech (Lake Forest, IL), 1M Tris pH 8.0 and 0.5 M EDTA from Quality Biologicals (Gaithersburg, MD), and isopropanol from Swan (Smyrna, TN). All other reagents for DNA purification were purchased from RPI Chemicals (Mount Prospect, IL). Magnetic rack bead separators for were purchased from EBay seller pochekailov (Seattle, WA). This seller offers an array of magnetic racks for various tube sizes that are, in our experience, the best available, and they are at a remarkably modest price point. Reagents for DNA quantitation, DNA gel stains, and qPCR were from Biotium (Fremont, CA). Oligonucleotides were obtained from IDT (Coralville, IA). Molecular biology reagents for post-purification quality assessment were from New England Biolabs (Ipswich, MA). Fluorescent quantitation of DNA was measured on a Qubit instrument (ThermoFisher, Waltham, MA). DNA gels were run using the electrophoresis apparatus from EmbiTech (San Diego, CA) and illuminated using a blue LED from IO Rodeo (Pasadena, CA). The optical filter for visualization of gel green stained gels was a 540 nm rapid edge filter from Omega optics (Austin, TX). Quantitative PCR was performed on a single channel open PCR machine from Chai (Santa Clara, CA).

## Methods

Two approaches were used to obtain the exact same chemical environments favorable for purification of cfDNA. Table 1 (“liquid-based method”) describes the reagents used in a solution-based approach that is convenient for small sample sizes. Magnetic beads are one of the most expensive components in the process and we found they are most effective when added in amounts proportional to volume.

**Table 1.** Liquid-based method for isolation of total DNA.

| Component | Composition | Relative volumes | 10 mL plasma prep | Cumulative volume | Notes |
| --- | --- | --- | --- | --- | --- |
| Plasma |  |  | 10 mLs | 10 mLs | Perform in 50 mL screw cap tube |
| Proteinase K, 20 mg/mL solution | 20 mg/mL Proteinase K, 50 mM Tris pH 8.0, 3 mM CaCl<sub>2</sub>, 50% glycerol v/v | Combine at a ratio of 1 volume Proteinase K solution per 50 volumes of plasma | 100 ul | 10.1 mLs | Mix prior to adding digestion buffer |
| Digestion buffer | 5M GITC, 25% Tween 20, 50 mM Tris pH 8.0, 25 mM EDTA | Combine at a ratio of ~2 volumes of digestion buffer per 3 volumes of plasma/Proteinase K | 6.5 mLs | 16.6 mLs | Heat to 56 C for ~ 1 hour |
| Binding buffer | 3.5 M GITC, 45% Isopropanol, 2.5% Tween 20, 10 mM Tris pH 8.0, 1 mM EDTA | Combine at a ratio of ~2 volumes of binding buffer to 1 volume of Plasma/Prot K/Digestion buffer | 33 mLs | 49.6 mLs | Mix prior to adding beads |
| 400 nM Silica beads | supplied as a 2.5 mg/mL solution | Combine at a ratio of 1 volume beads to 100 volumes of digested plasma in binding buffer | 400 ul | 50.0 mLs | Mix during addition of beads. Incubate 10 min at RT |
| Wash solution #1 | 3M GITC, 30% Isopropanol, 5% Tween 20, 40 mM Bis-Tris pH 6.0, 2 mM EDTA | For every 50 mL tube | 5 mLs |  | Perform in 5 mL tube |
| Wash solution #2 | 50 mM Tris pH 8.0, 0.5 mM EDTA, 80 % EtOH v/v | For every 50 mL tube | 5 mLs |  | Perform in 5 mL tube |
| 100% ethanol |  | For every 50 mL tube | 1 mL |  | Transfer to 1.5 mL tube. Aspirate and dry at 37 C |
| TE buffer | 10 mM Tris pH 8.0, 0.1 mM EDTA | For every 1.5 mL tube, elute with 100 ul then 60 ul | 100 ul, then 60 ul |  | Anticipate volume of ~150 ul of DNA. |

Therefore, to minimize costs for large volume purifications, we also devised a method in which pure and highly concentrated chemical constituents are added directly to plasma (Table 2; “solid-based method”).

**Table 2.**
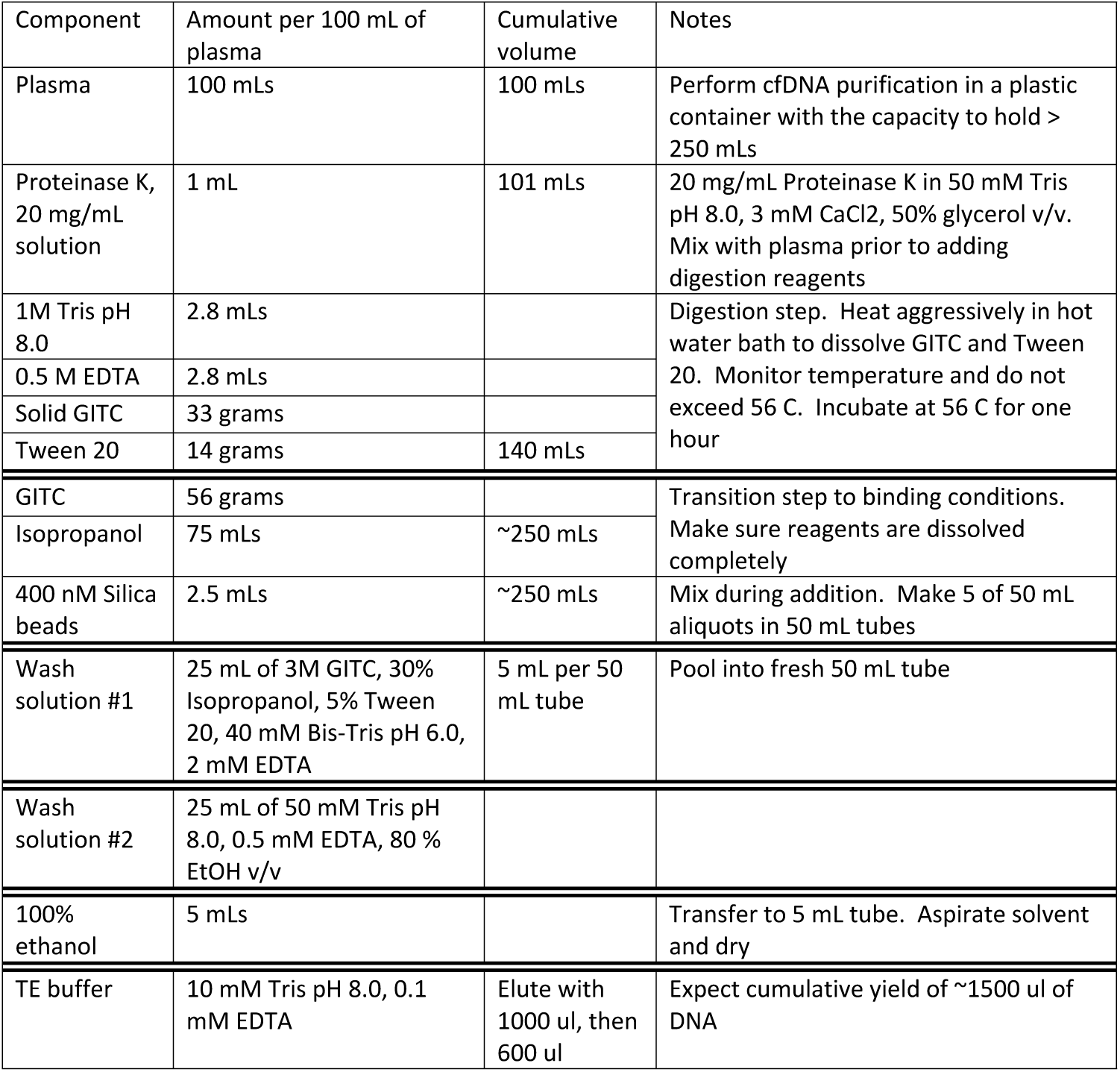
Solid-based method for isolation of total DNA.

Both methods need to be performed in plastic containers and not glass; glass is itself a silica surface that can bind DNA and drastically reduce yields. The first step is to combine proteinase K (formulated at 20 mg/ml in 50 mM Tris pH 8.0, 3 mM CaCl_2_ and 50% glycerol (v/v); store at 4° C) and plasma. Digestion reagents are then added. Reagents are dissolved by stirring and the reaction is brought to 56° C for approximately one hour. Binding reagents are then combined followed by the addition of beads. This slurry is brought to room temperature for about 5 min and then aliquoted into 50 mL conical centrifuge tubes. The tubes are placed in a magnetic separation rack. Once the beads are aggregated into a pellet, the supernatant is poured into a bio-hazard waste vessel. The beads are then washed with wash buffer #1, wash buffer #2, and 100% ethanol and dried completely. The total DNA is eluted with 15 ul of TE (10 mM Tris pH 8.0, 0.1 mM EDTA) per 1 mL of initial plasma input.

For size selection, combine 2 volumes of total DNA with one volume of DNA purification beads [13]. Incubate for 10 min at RT° C, pull aside beads, and transfer the 3 volumes of supernatant to a vessel containing 2 additional volumes of DNA purification beads. Incubate for 10 min, pull down beads (with bound cfDNA nucleosomal fragments), wash the bead pellet twice with an appropriate volume of 70% ethanol/water (v/v), and resuspend in 1 ul per 1 mL of plasma (the yield in ng/ul is also the original quantity in plasma in ng/mL).

For quality analysis, total DNA was purified using both the UltraPrep method and the QIAamp Circulating Nucleic Acid kit from Qiagen/Thermofisher (Hilden, Germany) as instructed by the manufacturer. RNAse A (Qiagen) treatment of QIAamp total DNA was performed by adding the enzyme directly to the total DNA (in elution buffer) to a final concentration of 10 ng/ul followed by incubation at 37° C for 30 min. The yield of total and size-fractionated DNA is measured using a Qubit fluorometer and AccuGreen™ reagents from Biotium. DNA gels are performed in 2% agarose with TBE buffer. The molecular size standards were the PCR marker from New England Biolabs which are 766, 500, 300, 150 and 50 bp. Alu quantitative PCR (qPCR) is performed with primers GAGGCTGAGGCAGGAGAATCG and GTCGCCCAGGCTGGAGTG [14] with OneTaq hot start (New England Biolabs) and EvaGreen™ dye (Biotium). The Cq values are converted into “Alu units” using the equation Alu units = power(10,-0.3*Cq+6) in Microsoft Excel. Library construction is evaluated by monitoring the attachment of adapters containing standard Illumina P5 and P7 sequences to cfDNA using a proprietary library construction technology (Ripple Biosolutions, Seattle, WA). The attachment efficiency is evaluated using PCR primers AATGATACGGCGACCACCGAGATCTACACTCTTTCCCTACACGACGCTCTTCCGATCT (Illumina-specific) and GAGGCTGAGGCAGGAGAATCG (Alu-specific) and qPCR conditions as described above. The results are quantified using a standard curve of premade cfDNA library material. The percent recovery for total DNA extraction and for size selection was determined by spiking this same library reference material into plasma prior to extraction or into total DNA prior to size selection. The percent recovery relative to the starting material was evaluated with the qPCR methods described above.

## Results

### cfDNA extraction and enrichment

The UltraPrep cfDNA purification method described here is a two-step process. The first step is extraction of total DNA from plasma with an emphasis on near-complete recovery of DNA present in the sample (Fig 1). The second step is size-based separation of nucleosomal-sized cfDNA fragments from high molecular weight genomic DNA. While the extraction technique described here is superficially similar to many other DNA extraction technologies, our method has several distinctive features. First, proteinase K is often the most expensive reagent used in DNA extraction procedures. To minimize cost, the conditions of the initial digestion step were configured to maximize proteinase activity and thereby allow reduced amounts of the enzyme to be used. This is described in more detail in supplemental S1 Fig. Second, several studies have shown that while DNA readily binds to silica surfaces in a variety of chemical conditions, a substantial fraction does so irreversibly. Here, the chemistry favors reversible association of DNA with silica beads and therefore robust recovery (∼84%) of total DNA (S2 Fig). Third, the DNA purification method is completely passive and therefore mechanical devices such as vacuum pumps or centrifuges are not required. Fourth, the method is scalable, meaning the yield of extracted DNA per milliliter of input is consistent across a broad range of sample input volumes.

**Fig 1.**
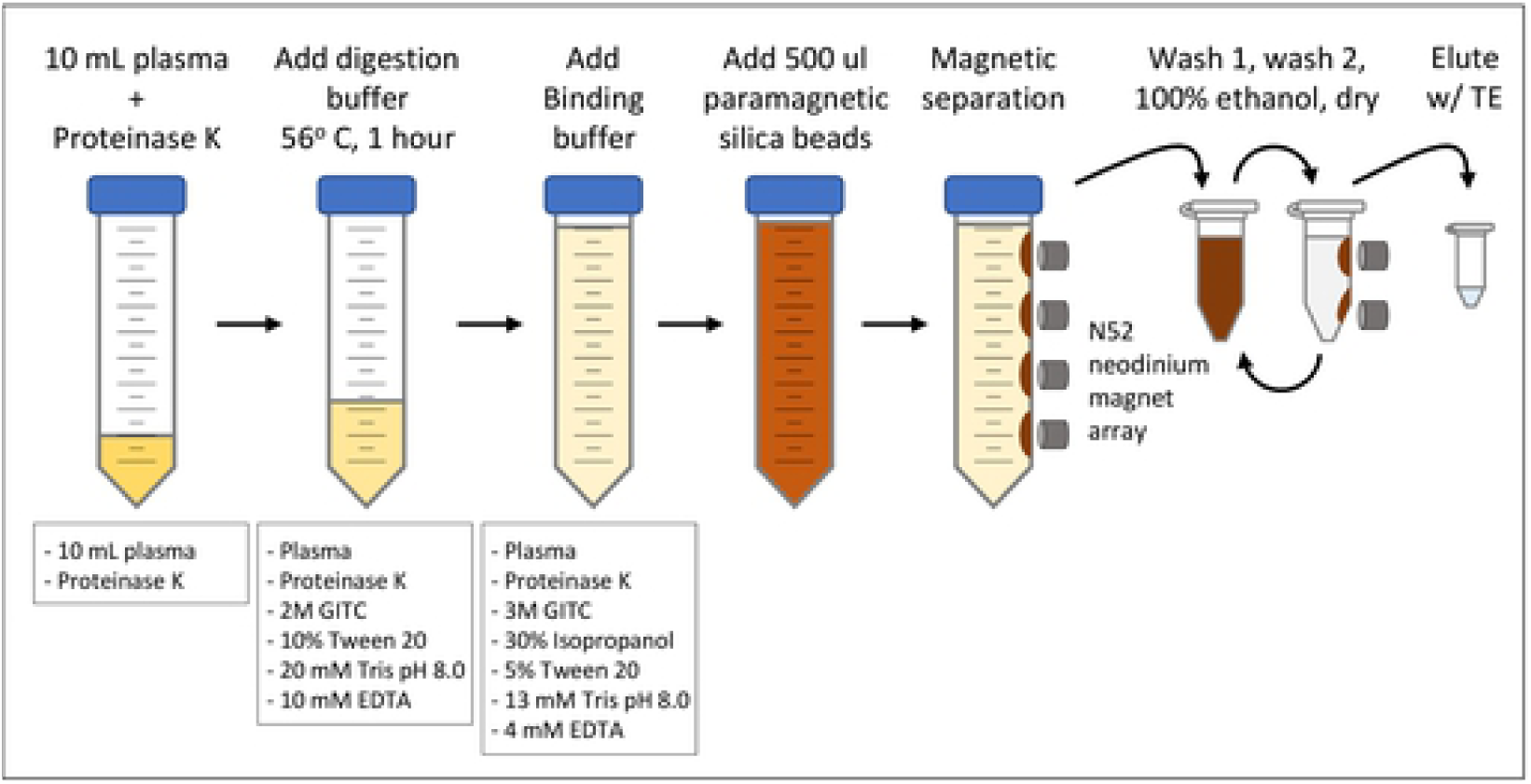
UltraPrep procedure for purification of total DNA from plasma using the liquid-based method.

We developed two related approaches for DNA extraction. For smaller sample volumes, a liquid-based protocol that utilizes additions of premade buffers is outlined in Table 1. This is the method illustrated schematically in Fig 1. For larger samples, we developed a method that involves additions of concentrated materials directly to plasma in amounts that recapitulate the chemical environment most favorable for high yield recovery of DNA (Table 2). The former approach is convenient while the latter strategy minimizes cumulative sample volume and therefore the amount of somewhat costly silica beads needed to fully recover total DNA (S3 Fig).

Total DNA extracted from plasma is most often a mixture of nucleosomal fragments and high molecular weight genomic DNA. In the method presented here, these two species can be partitioned into separate fractions using a two-step bead purification that is commonly referred to as “double-sided solid phase reversible immobilization (SPRI)” (Fig 2). Polyethylene glycol (PEG), and to a lesser extent salt, drive binding of DNA onto the surface of carboxyl-coated magnetic beads. The core principle behind double-sided size separation is that there is an inverse relationship between the concentration of PEG and the size of bound DNA fragments. In the first step a more dilute concentration of PEG favors binding of high molecular weight DNA. The supernatant is then added to additional PEG (and SPRI beads) in the second step to recover the nucleosomal cfDNA fragments. The overall recovery of DNA from this enrichment step was about 80% (S4 Fig). This generates an estimate that about 2/3 of the nucleosomal cfDNA fraction (84% from step1 × 80% from step 2 = 67% overall) was recovered in the UltraPrep process. Coincidentally, in the plasmapheresis samples we have worked with, about 1/3 of the total DNA is high molecular weight and 2/3 is nucleosomal cfDNA (for example, see Fig 2).

**Fig 2.**
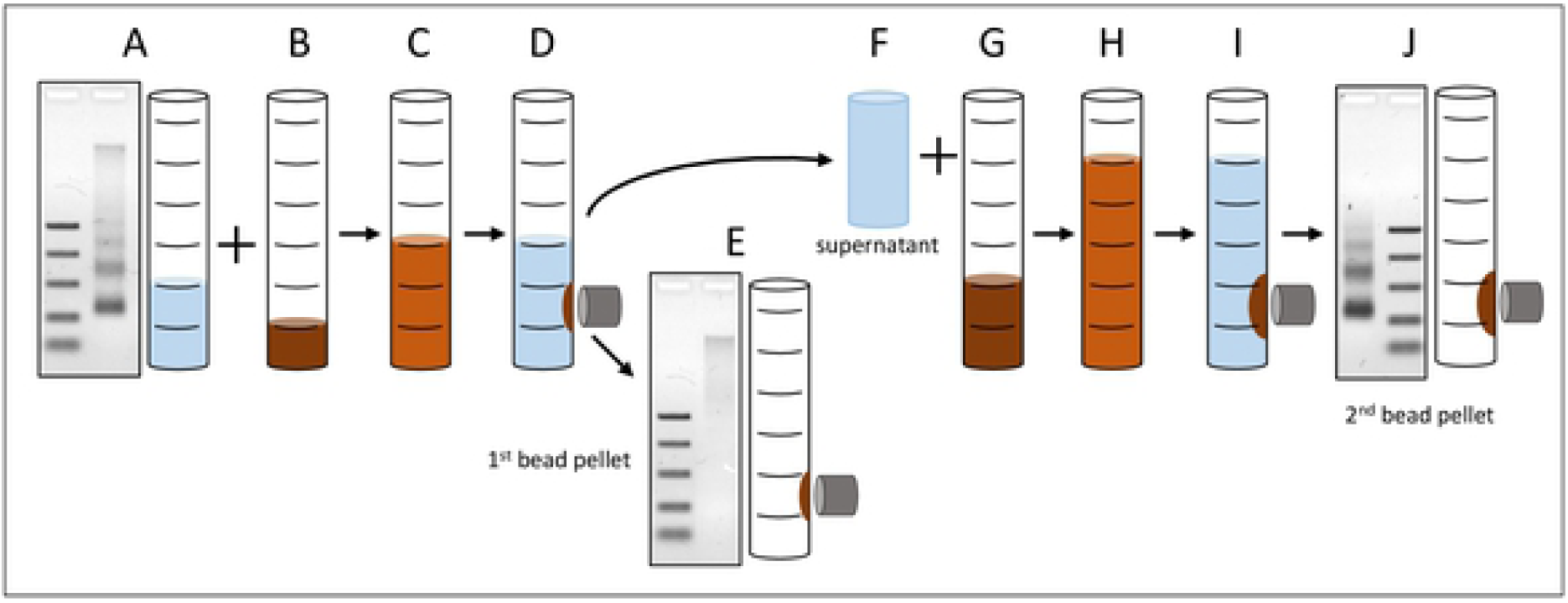
Double-sided SPRI bead enrichment of nucleosomal cfDNA fragments. (A) Two volumes of total DNA from the first stage of the UltraPrep procedure are (B) combined with one volume of DNA purification beads [13]. (C) After a 10 min incubation at RT° C, (D) the beads are pulled aside and the (F) three volumes of supernatant are transferred to (G) two additional volumes of DNA purification beads. (H) The blended mixture is incubated for 5 min and (I) the beads with bound cfDNA are pulled aside, washed with 70% ethanol/water, dried and (J) the DNA is eluted with TE.

### Performance

With an eye toward both research applications and clinical utilization of the cfDNA purified using the UltraPrep method, we established four independent assays and a comnparison with the industry-standard method to evaluate UltraPrep purified material (Fig 3). First, we measured the yield of double-stranded DNA (dsDNA) using dsDNA-specific fluorescent dyes and a Qubit fluorometer. The typical yield of purified, nucleosomal-sized cfDNA from healthy donor plasmas collected by automated plasmapheresis into sodium citrate was 3 – 4 ng per mL of plasma (S1 Table). Second, we used agarose gel electrophoresis to determine size distribution of purified material. Acceptable samples exhibited a fragmentation pattern consistent with DNAse cleavage in the linker region between adjacent nucleosomes. Third, several downstream analytical techniques (e.g. ddPCR, targeted amplicon sequencing, BEAMing, etc.) require that the input cfDNA is a robust amplification template devoid of inhibitors. We used qPCR measurement of human Alu sequences to establish that the specific “amplifiability” per picogram of purified cfDNA was consistent between purified lots of material and similar to cfDNA purified using the industry standard (QIAamp). Fourth, quantitative NGS methods for cfDNA analysis rely on the attachment of adapter sequences as a prerequisite for creating genomic cfDNA libraries. We monitored the “clonability” of purified cfDNA by measuring the attachment efficiency of adapters containing standard Illumina sequences to cfDNA. This measurement was performed by qPCR with a primer pair where one primer was specific for the adapter sequence and the other was specific for the human Alu repeat. Using standard curve analysis with an established control, the readout of nanograms of adapter-modified cfDNA per nanogram of input was a direct measurement of adapter attachment efficiency. Finally, most published studies cite the QIAamp Circulating Nucleic Acid kit from Qiagen as the method used for initial purification of cfDNA; in other words, this is the established purification technology by which other methods should be benchmarked. We routinely compared the assay performance metrics for cfDNA purified from the same plasma using the QIAamp procedure and the UltraPrep method.

**Fig 3.**
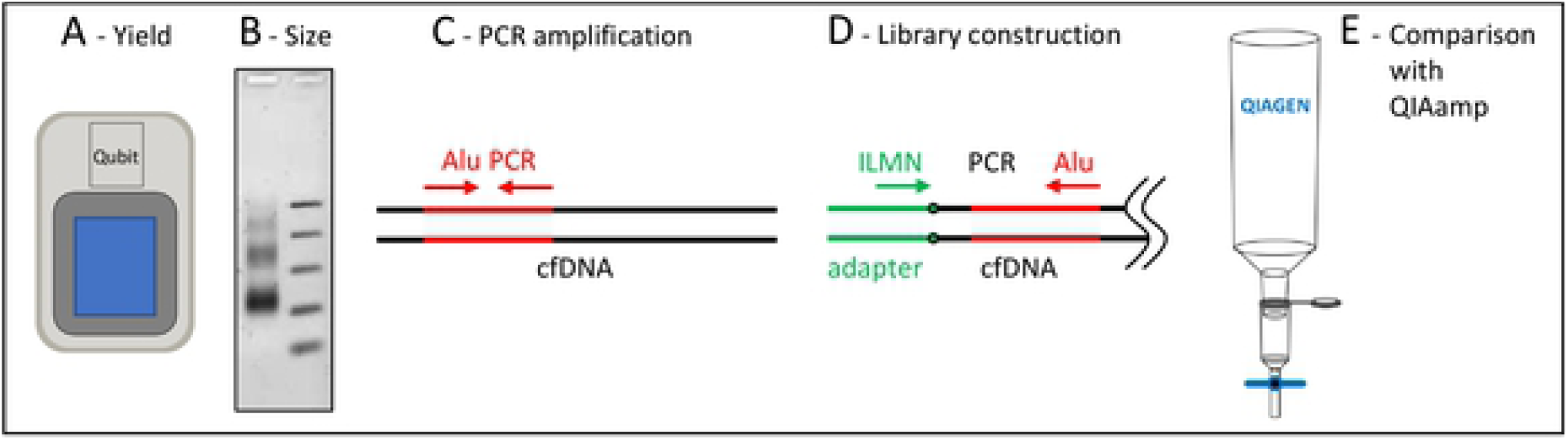
Four quality assays and one comparison used to evaluate purified cfDNA. (A) The Qubit fluorometer was used to quantify the amount of double-strand-DNA-specific dye bound to DNA. (B) Agarose gel electrophoresis was used to assess the fragmentation pattern of purified cfDNA. (C) Alu sequence-specific qPCR with primers directed to the human Alu sequence [14] were used to determine the amplifiability of cfDNA with a readout of Alu units detected/pg of DNA. (D) Library construction efficiency was determined by qPCR as the percentage of cfDNA ends attached to an adapter that contains standard Illumina NGS sequences. (E) An aliquot of the plasma samples used in large scale preparations was purified using the industry standard QIAamp technology, and the resulting DNA from both methods was compared using the quality assays described in (A) through (D).

The comparisons with QIAamp purified cfDNA merit further consideration (Fig. 4). Our initial observation was that QIAamp-purified cfDNA had less specific activity for Alu content and lower rates of adapter attachment than DNA purified by the method described here. Further investigation revealed two reasons for this. First, the QIAamp kit is a total DNA isolation method that collects both high molecular weight genomic DNA and cfDNA fragments. High molecular weight DNA, present to some extent in many plasma samples, performs poorly in the library construction assay, thereby accounting for some of the discrepancy. Second, we found that the carrier RNA routinely added during QIAamp purification is a significant interfering substance. It falsely elevates Qubit readings, resulting in overestimation of DNA concentrations by as much as 50%. In the example shown in Fig. 4, the initial total DNA Qubit reading for the QIAamp extracted sample indicated a yield of 12.2 ng/mL plasma. After RNAse A treatment (see Methods), this value dropped to 8.3 ng/mL plasma. Using the same lot of plasma, the total DNA yield from the UltraPrep protocol that does not use carrier RNA was a comparable value of 9.3 ng/mL plasma. After size selection, the yield of nucleosomal fragments from the two methods was essentially the same (Fig. 4A). Similarly, the size distribution, Alu PCR and library construction results were more or less identical for both sets of samples.

**Fig. 4.**
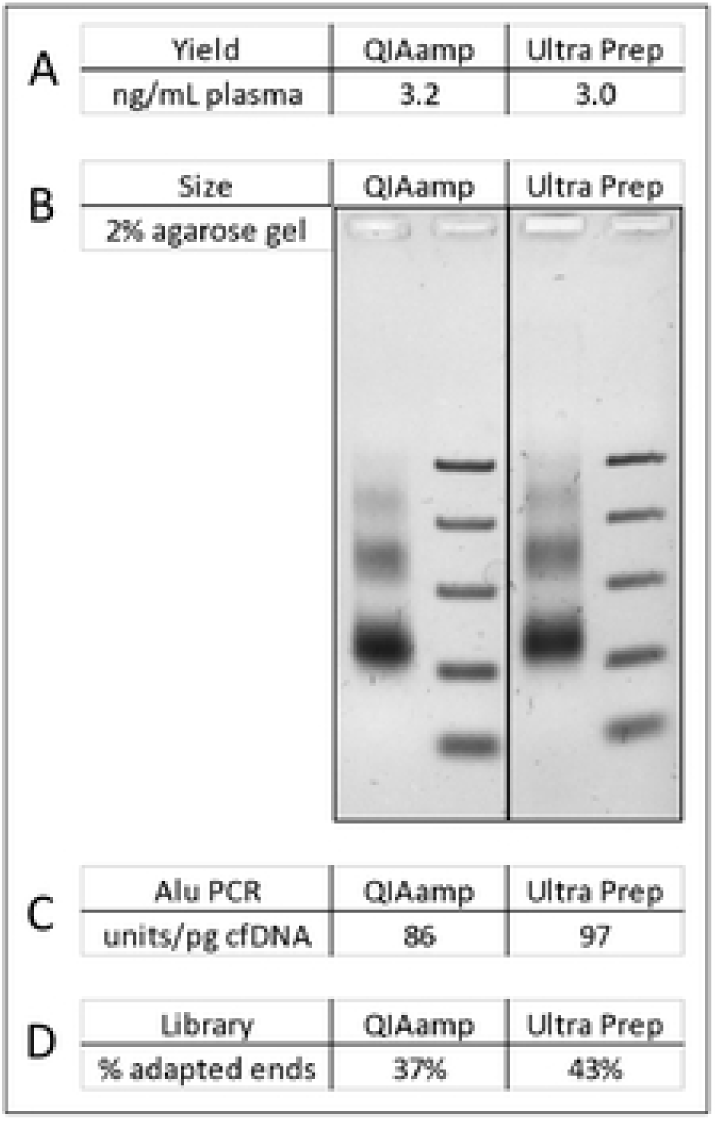
Quality analysis of nucleosomal sized DNA purified from a QIAamp prep and from UltraPrep total DNA. (A) Total yield of cfDNA fragments. (B) Size distribution of cfDNA. (C) Comparison of Alu units per pg. (D) Comparison of adapter attachment efficiency.

The UltraPrep method was also successfully applied to whole blood samples collected in lavender-top K_2_EDTA vacutainer tubes (S1 Table). The yield of nucleosomal-sized cfDNA was rather high in these samples, which, based on equivalent yields from QIAamp and on previous studies [12], we believe to be a characteristic of the sample and not the collection method. The method was also applied to unspun urine that was collected in EDTA-containing vessels. Most of the resulting DNA was high molecular weight, with a broad smear present in the low molecular size fraction (data not shown).

## Discussion

The UltraPrep open-source method for purification of cfDNA represents a significant advance in the ability to access this vital diagnostic analyte. It represents a very significant reduction in cost from currently used methods. The cost of cfDNA isolation from human plasma using the current industry standard QiaAmp technology is approximately five dollars per mL of plasma processed. The total cost of reagents and consumables using the UltraPure process is approximately 50 cents per mL of plasma processed. The yield (ng per mL plasma) of purified nucleosomal fragments from the two methods is indistinguishable. The UltraPrep protocol scales easily from a few mLs of plasma to hundreds of mLs of plasma with little change in the time and effort required for cfDNA purification. Small- and large-scale purifications can easily be completed in a single day. The resulting purified material performs exceptionally well in downstream analytical assays. The size selection step addresses a major preanalytical concern by removing high molecular weight genomic DNA that can introduce artifacts into downstream analysis, especially those methods that are attempting to measure the minor allele frequency of rare tumor markers. In our view, this method is preferable to using fixative-containing blood collection tubes to stabilize blood cells. The same reagents that mitigate cell breakage can potentially cross-link DNA and thereby confound test results.

The UltraPrep method makes purification of microgram quantities of cfDNA from single-donor plasmapheresis collections feasible. This in turn opens new opportunities. For instance, the same cfDNA sample can conceivably be used for diagnostic research, assay development, and testing implementation. A panel of donor samples can be used time and again to calibrate the background noise in newly developed genomic assays. This is particularly important in the case of next-generation sequencing applications where systematic error can generate false positive signals. Moreover, there is an acute need for “truth samples”, comprised of *bona fide* cfDNA, that can be used for proficiency testing. The current paradigm of comparing cancer patient cfDNA with matched DNA extracted from tumor biopsies invariably generates discrepancies that are most often explained away as biological phenomenon [15]. Similarly, “synthetic cfDNA” spiked with known markers is an uncertain approximation of genuine, physiologically generated DNA [12]. Rather we propose that proficiency testing can feasibly be accomplished by monitoring common genetic polymorphisms in systematically blended cfDNA samples from two unrelated donors [4]. Lastly, our overarching goal is to see tests for early detection of cancer that are conducted during routine wellness exams. Most asymptomatic individuals are capable of donating the quantities of whole blood that will be needed for deep genomic coverage detection tests. UltraPrep technology has the scale to accommodate these higher volume plasma samples.

## Supporting information

**S1 Fig. Yield of total DNA as a function of proteinase K addition.** Identical 10 mL aliquots of several different donor samples were processed using concentrations of proteinase K shown. The quantity of 200 ug/mL plasma (10 ul of a standard 20 mg/ml solution of enzyme added per mL of plasma) that was chosen for the protocol is highlighted in green.

**S2 Fig. Recovery of spiked-in DNA from four replicates of UltraPrep.** The spike-in material was completed cfDNA library with Illumina adapter sequences. One ng per mL of plasma was added. The amount of library in the preprocessed control and four replicates was determined by qPCR using the Illumina + Alu primer pair. The average recovery from the four samples was 84%.

**S3 Fig. Yield of total DNA as a function of silica bead concentration.** Identical 10 mL aliquots of plasma were processed using the microliters of beads/prep volume shown. The yield of total DNA after the initial purification step and of high molecular weight versus nucleosomal-sized DNA is after the size selection step are shown as a function of added bead volume (2.5 mg/mL beads). The prep volume per mL of input plasma is larger for the liquid-based prep than for the solid-based prep by a factor of two-fold. In consideration of this, the quantity of 8 ul/mL prep volume was chosen for the liquid-based prep and 10 ul/mL prep volume for the solid-based prep.

**S4 Fig. Recovery of spiked-in DNA after size fractionation.** The spike-in material was completed cfDNA library with Illumina adapter sequences. One ng per mL of plasma was added. The amount of material eluted from the first and second bead pellets was determined by qPCR using the Illumina + Alu primer pair. “Total” recovery is the sum of the two elutions.

**S1 Table. Yields of cfDNA across time and samples.**

